# HLA-B*39:06 Efficiently Mediates Type 1 Diabetes in a Mouse Model Incorporating Reduced Thymic Insulin Expression^1^

**DOI:** 10.1101/264531

**Authors:** Jennifer Schloss, Riyasat Ali, Jeremy J. Racine, Harold D. Chapman, David V. Serreze, Teresa P. DiLorenzo

**Affiliations:** Department of Microbiology and Immunology, Albert Einstein College of Medicine, Bronx, NY 10461; The Jackson Laboratory, Bar Harbor, ME 04609; Department of Medicine (Division of Endocrinology), Albert Einstein College of Medicine, Bronx, NY 10461

## Abstract

Type 1 diabetes (T1D) is characterized by T cell-mediated destruction of the insulin-producing βcells of the pancreatic islets. Among the loci associated with T1D risk, those most predisposing are found in the MHC region. HLA-B*39:06 is the most predisposing class I MHC allele and is associated with an early age of onset. To establish an NOD mouse model for the study of HLA-B*39:06, we expressed it in the absence of murine class I MHC. HLA-B*39:06 was able to mediate the development of CD8 T cells, support lymphocytic infiltration of the islets, and confer T1D susceptibility. Because reduced thymic insulin expression is associated with increased T1D risk in patients, we incorporated this in our model as well, finding that HLA-B*39:06-transgenic NOD mice with reduced thymic insulin expression have an earlier age of disease onset and a higher overall prevalence as compared to littermates with typical thymic insulin expression. This was despite virtually indistinguishable blood insulin levels, T cell subset percentages, and TCR Vβ family usage, indicating that reduced thymic insulin expression does not impact T cell development on a global scale. Rather, we propose that it allows the thymic escape of insulin-reactive HLA-B*39:06-restricted T cells which participate in β cell destruction. We also found that in mice expressing either HLA-B*39:06 or HLA-A*02:01 in the absence of murine class I MHC, HLA transgene identity alters TCR Vβ usage, which may contribute to varying diabetogenic CD8 T cell repertoires in the presence of different HLA class I alleles.

## INTRODUCTION

Type 1 diabetes (T1D)^3^ is characterized by T cell-mediated destruction of insulin-producing β cells (1). Both CD4 and CD8 T cells are important for T1D pathogenesis, with CD8 T cells requiring the presentation of β cell epitopes by class I MHC molecules in order to interact with, and eliminate, the β cells (2, 3). It is thus unsurprising that while multiple genetic loci have been found to contribute to T1D development, those most predisposing to T1D can be found in the MHC region (4). Several class I MHC alleles have been found to be predisposing to T1D, including HLA-A*02:01 and HLA-B*39:06 (5-9). While the presentation of β cell epitopes by HLA-A*02:01 has long been known and extensively studied (10), HLA-B*39:06 has only more recently gained attention as a T1D-associated allele, and much remains to be understood about its ability to confer T1D risk.

While T1D associations have been observed at all HLA class I loci (9), HLA-B*39:06 is the most predisposing HLA class I allele (7, 9) and, importantly, is associated with an early age of onset (11). Furthermore, HLA-B*39:06 is most common among the Latin American population (12), where T1D incidence has been rising (13-15). Development of an HLA-B*39:06-transgenic mouse model is thus of the utmost importance in order to understand the relationship between HLA-B*39:06, genetic risk background, and T1D pathogenesis. A transgenic model is also essential for the preclinical testing of HLA-B*39:06-targeted treatments.

Given the multiple risk factors associated with T1D predisposition, it is important to study HLA-B*39:06 in a translationally relevant manner. The NOD mouse is considered by many to be a good model for human T1D (16, 17). For example, the NOD class II MHC H2-A^g7^ shares striking similarities with several T1D-associated human class II MHC alleles such as HLA-DQ8 (18). Among other similarities, both NOD mice and human T1D patients display reduced regulatory T cell function and reduced IL-2 signaling (17, 19, 20). Most importantly, T cells from HLA-transgenic NOD mice may target similar or even identical β cell epitopes to those found in T1D patients (21-23). However, to most accurately model HLA-B*39:06 in the context of human T1D, it is preferable to incorporate additional human non-MHC risk alleles. In humans, the non-MHC locus that confers the most susceptibility to T1D is the variable number of tandem repeats (VNTR) region of the insulin gene (24-26). Shorter VNTR sequences are known as class I while longer VNTR sequences are known as class III. Class I VNTR sequences are associated with T1D risk and with a decrease in thymic insulin mRNA levels compared with the longer class III VNTR alleles, which are protective (24). With a resultant decrease in thymic insulin expression, there is a hypothesized increase in the escape of insulin-reactive T cells. This may explain the association between class I insulin VNTR alleles and T1D predisposition in humans (25).

The reduced thymic insulin expression seen in T1D patients may be modeled in mice through introduction of one or two *Insulin 2* (*Ins2*) knockout (KO) alleles (27-31). Mice possess two insulin genes, *Insulin 1* (*Ins1*) and *Ins2*. Although expressed in the pancreas, little (28, 32) to no (33, 34) *Ins1* expression occurs in the thymus. In contrast, *Ins2* is expressed in both the thymus and the pancreas (28), and both NOD mice and mouse strains not prone to T1D exhibit altered T cell tolerance to insulin upon *Ins2* ablation (28, 30, 31, 35). We have shown that NOD mice even just heterozygous (Het) for the *Ins2*^KO^ allele exhibit decreased thymic insulin expression as seen in human T1D patients (27). In the context of HLA-A*02:01, we have previously found that NOD mice with reduced thymic insulin expression display increased T1D incidence, islet infiltration, and CD8 T cell responses to insulin (27, 29), speaking to the importance of examining multiple risk alleles simultaneously.

Here we have developed HLA-B*39:06-transgenic NOD mouse models and have demonstrated that HLA-B*39:06 is able to independently mediate the development of CD8 T cells required for T1D onset. In the context of reduced thymic insulin expression, HLA-B*39:06-transgenic NOD mice develop T1D at an accelerated rate compared to mice with wild-type (WT) thymic insulin expression, despite normal blood insulin levels and no gross alterations in lymphocyte composition or TCR Vβ family usage. We propose that with a decrease in thymic insulin expression, HLA-B*39:06 is less able to negatively select insulin-specific CD8 T cells and, with the high concentration of insulin found in the islets, is able to present insulin epitopes to escaped CD8 T cells. Thus, by generating HLA-B*39:06-transgenic NOD mice in the presence of reduced thymic insulin expression, we show here the development of models that will provide excellent tools for the examination of HLA-B*39:06’s impact on T1D and for the preclinical testing of HLA-B*39:06-targeted therapies.

## MATERIALS AND METHODS

### Mice

To develop HLA-B*39:06-transgenic NOD mice, we prepared a monochain chimeric HLA-B*39:06 construct, comprising the α1 and α2 peptide binding domains of HLA-B*39:06 linked to the α3 CD8 binding and transmembrane domains of H2-D^b^ with human β_2-_microglobulin (β2m) linked covalently to the α1 domain. Chimeric constructs of this design are designated human β2m/HLA/H2-D^b^ (HHD) (36). This HLA-B*39:06 HHD construct was injected into NOD zygotes, and founder mice were identified by PCR of tail-tip DNA using these HLA-B*39:06 primers: 5′-CTTCATCTCAGTGGGCTAC-3′ and 5′-CGGTCAGTCTGTGTGTTGG-3′. Positive progeny were further assessed for HLA-B*39:06 expression on their peripheral blood leukocytes by flow cytometry using anti-HLA-A, B, C (W6/32; BioLegend). Founder 45, with the highest expression of HLA-B*39:06, was selected for further investigation and was crossed with an NOD mouse. Progeny of this cross were assessed for the presence of the transgene by PCR of tail-tip DNA; mice hemizygous for the transgene were designated NOD.HLA-B*39:06^Hemi^. To maintain this strain, NOD.HLA-B*39:06^Hemi^ mice were crossed with NOD littermates. NOD.HLA-B*39:06^Hemi^ females were also crossed with male mice from NOD.β2m^KO^ (37) or NOD.β2m^KO^. Ins2^KO^ strains (27). To fix the HLA-B*39:06 transgene to homozygosity, the resulting progeny were interbred as appropriate to generate HLA-B*39:06 homozygous mice (HLA-B*39:06^Hom^) with the following genotypes: NOD.HLA-B*39:06^Hom^.β2m^KO^, NOD.HLA-B*39:06^Hom^.β2m^KO^.Ins2^Het^, and NOD.HLA-B*39:06^Hom^.β2m^KO^.Ins2^KO^. As we and others have found that female NOD.β2m^KO^ mice breed poorly (38), we crossed male β2m^KO^ mice with female β2m^Het^ mice whenever possible. The WT and KO β2m and WT and KO Ins2 alleles were identified by PCR using the following primer pairs: β2m^WT^: 5′-GAAACCCCTCAAATTCAAGTATACTCA-3′ and 5′-GACGGTCTTGGGCTCGGCCATACT-3′; β2m^KO^: 5′-GAAACCCCTCAAATTCAAGTATACTCA-3′ and 5′-TCGAATTCGCCAATGACAAGACGCT-3′; Ins2^WT^: 5′-GGCAGAGAGGAGGTGCTTTG-3′ and 5′-AGAAAACCACCAGGGTAGTTAGC-3′; Ins2^KO^: 5′-GGCAGAGAGGAGGTGCTTTG-3′ and 5′-ATTGACCGTAATGGGATAGG-3′. NOD.HLA-A*02:01 (HHD).β2m^KO^ mice have been previously described (21).

### Assessment of HLA-B*39:06 homozygosity by real-time PCR

Mouse tails were numbed with ethyl chloride (Gebauer) and the tail tips were removed. Tails were digested in 200 µl proteinase K (Roche) solution overnight at 56°C. The reaction was stopped by placing tails at 95°C for 10 min. The resultant DNA (1 µl) was mixed with PrimeTime Gene Expression Master Mix (IDT) and each of the following primers and TaqMan probes: HLA-B*39:06 primers (5′-TTCATCTCAGTGGGCTACG-3′ and 5′-TGTGTTCCGGTCCCAATATTC-3′) and probe [5′-(6-FAM)-TCGCTGTCGAACCTCACGAACTG-(Zen probe with Iowa Black)-3′]; internal positive control primers (5′-CACGTGGGCTCCAGCATT-3′ and 5′-TCACCAGTCATTTCTGCCTTTG-3′) and probe [5′-(Cy5)-CCAATGGTCGGGCACTGCTCAA-(Black Hole Quencher 2)-3′]. Real-time quantitative PCR was performed in triplicate using an iQ5 Real-time PCR Detection System (Bio-Rad). Amplification was carried out as follows: initial denaturing at 94°C for 2 min, followed by 38 cycles of 20 s at 94°C, 15 s at 60°C and 10 s at 72°C. Copy numbers were calculated using the 2^ΔΔCt^ method.

### Assessment of T1D

Mice were monitored weekly from 4-30 wks for glucosuria using Diastix reagent strips (Bayer). Mice were considered diabetic following two consecutive positive tests. The first positive test was recorded as the date of diabetes onset.

### Histology

Pancreata were fixed in Bouin’s solution, sectioned at three non-overlapping levels, and stained with aldehyde fuchsin and hematoxylin and eosin. Islets were scored for insulitis by a blinded observer as previously described (39): 0, no visible lesions; 1, peri-insular or non-invasive leukocytic aggregates; 2, <25% islet destruction; 3, 25-75% islet destruction; 4, >75% islet destruction. A mean insulitis score was determined for each mouse by dividing the total score for each pancreas by the total number of islets examined. Diabetic mice were assigned a score of 4.

### Blood collection and staining of peripheral blood leukocytes

Blood (10 µl) was collected from the mouse tail vein and added to 50 µl PBS (pH 7.2, Gibco) with 1 mM EDTA (Sigma). Samples were mixed well and erythrocytes were lysed for 2-3 min with 200 µl ACK lysis buffer (Lonza). Plates were centrifuged at 700xg for 3 min and ACK lysis was repeated. Following centrifugation, samples were washed twice with PBS containing 1% FBS (HyClone) and 0.1% (w/v) sodium azide. All subsequent washes and dilutions were performed using this buffer. Cells were stained with Fc Block (BD Biosciences), followed by anti-CD8α (53-6.7; BD Biosciences) and anti-HLA-A, B, C (B9.12.1; Beckman Coulter) and incubated on ice for 15-20 min. Samples were washed twice, suspended in 1 μg/ml DAPI and incubated on ice for 15-30 min. Samples were filtered through a 35-µm cell strainer prior to data collection on a BD LSRII flow cytometer with five lasers (355 nm, 405 nm, 488 nm, 561 nm and 640 nm). Data were analyzed using FlowJo software (version 8.8.6).

### Serum collection and insulin ELISA

Blood (20-40 µl) was collected from the mouse tail vein and allowed to clot at room temperature for 1 h. Samples were centrifuged for 15 min at 960xg at 4°C. Serum was stored in aliquots at −20°C. Blood insulin levels were measured using the Mouse Ultrasensitive Insulin ELISA (ALPCO). Absorbance of each well at 405 nm was detected using an Emax precision microplate reader (Molecular Devices) and the results were analyzed using GraphPad Prism 7 software.

### Splenocyte preparation and flow cytometry

Mice were euthanized using CO_2_ asphyxiation, followed by cervical dislocation. Spleens were harvested and placed in ice-cold RPMI (Gibco) supplemented with 10% FBS, 1% sodium pyruvate (Gibco), 1% non-essential amino acids (Gibco), 50 U/ml penicillin and 50 µg/ml streptomycin (Gibco). Spleens were crushed, passed through a 40-µm cell strainer and washed with RPMI. Samples were centrifuged at 486xg for 5 min. Erythrocytes were lysed in ACK lysis buffer (Lonza) for 4 min at room temperature and washed with RPMI. The resultant cells were centrifuged and washed twice with PBS. Prior to the final wash, samples were passed through a 40-µm cell strainer. Cells were counted and suspended in PBS. Samples prepared in the above manner were added to a V-bottomed plate and centrifuged at 486xg for 5 min. Samples were washed once in PBS containing 2% FBS (HyClone). This buffer was used for all subsequent washing and dilution steps. Cells were stained with Fc Block (BD Biosciences) on ice for 10 min and washed once. For monitoring of class I MHC expression, cells were incubated on ice for 20 min with labeled anti-HLA-A, B, C (B9.12.1; Beckman Coulter), anti-pan murine class I MHC (M1/42; The Jackson Laboratory), or an appropriate isotype control antibody (mouse IgG2a for B9.12.1 and rat IgG2a/κ for M1/42). For analysis of splenic immune cell populations, cells were stained with labeled anti-CD19 (6D5; BioLegend), anti-TCRβ (H57-597; BD Biosciences), anti-CD8α (53-6.7; BD Biosciences), anti-CD4 (GK1.5; BD Biosciences), and anti-CD25 (PC61.5; eBioscience). For study of TCR Vβ usage, an anti-mouse TCR Vβ screening panel was used (BD Biosciences) in conjunction with labeled anti-CD19 (6D5; BioLegend), anti-CD3ε (145-2C11; BD Biosciences), anti-CD8α (53-6.7; BD Biosciences), anti-CD4 (GK1.5; BD Biosciences), and anti-CD25 (PC61.5; eBioscience). Samples were washed twice, incubated in 1 μg/ml DAPI for 15 min on ice, and filtered through a 35-µm cell strainer prior to data collection. Data were collected on a BD LSRII flow cytometer with five lasers (355 nm, 405 nm, 488 nm, 561 nm and 640 nm) and analyzed using FlowJo (version 8.8.6) and GraphPad Prism 7 software.

## RESULTS

### NOD mice transgenic for HLA-B*39:06 are susceptible to T1D

To begin to study the association of HLA-B*39:06 with T1D, we first developed NOD.HLA-B*39:06 mice using a monochain HLA-B*39:06 construct. We tracked these mice for susceptibility to disease to ensure that the integration of HLA-B*39:06 did not interfere with T1D development. We found no decrease in disease susceptibility compared to non-transgenic littermates in either females (Fig. 1A) or males (Fig. 1B). The earliest age of onset among female mice was 14 wks, with 82% diabetic by 30 wks. In males, the earliest age of onset was at 13 wks, though as expected, incidence was reduced compared to females, with only 57% converting to disease by 30 wks of age. Because females were more susceptible to disease than males, we used female mice for our subsequent experiments.

**Figure 1.**
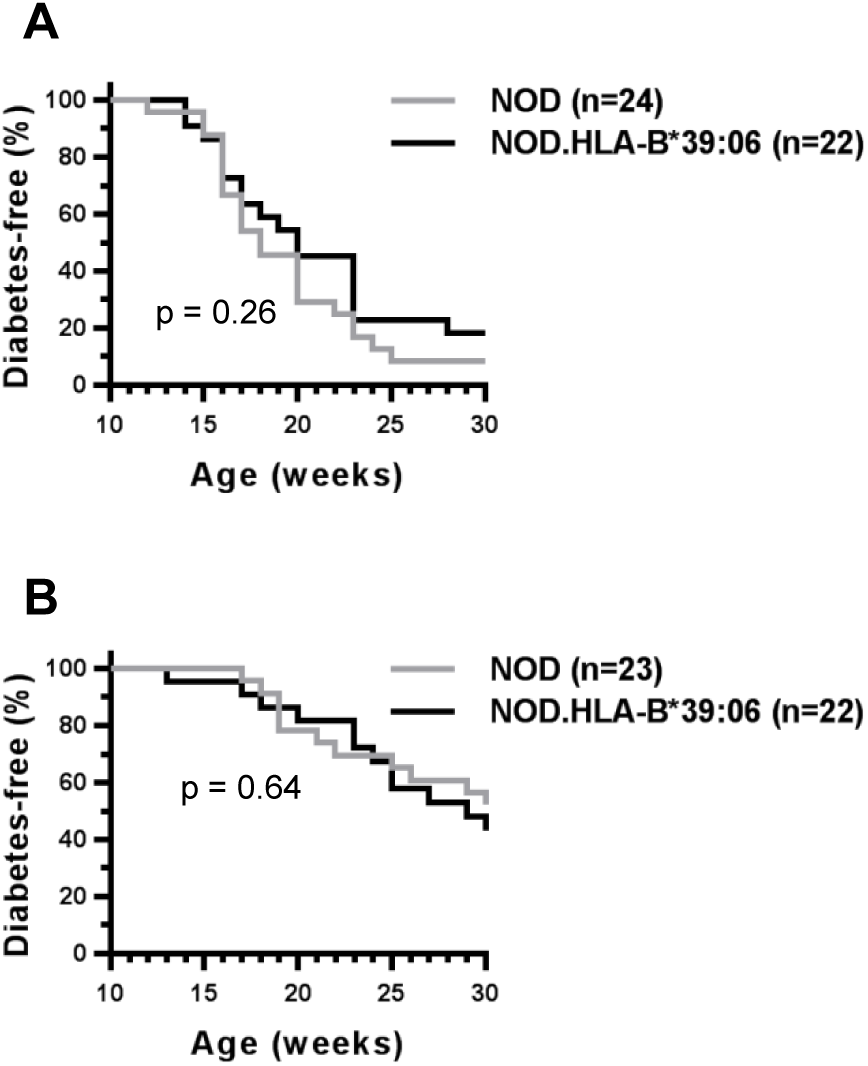
HLA-B*39:06-transgenic NOD mice are susceptible to T1D. Diabetes incidence curves for female (**A**) and male (**B**) NOD.HLA-B*39:06 mice and non-transgenic NOD littermates are shown. (A) p = 0.26, Mantel-Cox; (B) p = 0.64, Mantel-Cox.

### HLA-B*39:06 allows for the selection of CD8 T cells in NOD mice

To examine the influence of HLA-B*39:06 on T1D without the complicating factor of the concomitant expression of murine class I MHC molecules, we developed a model in which the transgenic HLA-B*39:06 was expressed in the absence of murine β2m by breeding with the NOD.β2m^KO^ strain (37). Because the transgenic HLA-B*39:06 HHD molecules contain covalently bound human β2m, HLA-B*39:06 can fold without reliance on murine β2m, whereas the endogenous H2-K^d^ and H2-D^b^ cannot. To maximize the expression of HLA-B*39:06 and the thymic selection of CD8 T cells, we sought to fix the HLA-B*39:06 transgene to homozygosity (HLA-B*39:06^Hom^). To do this, we first examined the level of human class I MHC on peripheral blood leukocytes from female NOD.HLA-B*39:06.β2m^KO^ mice (Fig. 2A). While all mice tested expressed human class I MHC, there appeared to be two groups of mice, one with high levels of class I MHC, with an average geometric mean fluorescence intensity (MFI) of 1534, and one with low class I MHC levels, with an average MFI of 607, suggesting that the mice with increased human class I MHC levels were HLA-B*39:06^Hom^. We used real-time PCR for the HLA-B*39:06 transgene to ensure that the high expressers were indeed homozygous for HLA-B*39:06 (Fig. 2B). We found that the average copy number of the low expressers was 1.6. This value was consistent with previous experiments with NOD.HLA-B*39:06.β2m^KO^ mice that were known to be hemizygous (data not shown), confirming that the human class I MHC-low mice were HLA-B*39:06^Hemi^. Human class I MHC-high mice had a copy number of 3.1, nearly double what was seen in the HLA-B*39:06^Hemi^ mice and indicating that these mice were, in fact, homozygous for HLA-B*39:06. We hypothesized that HLA-B*39:06^Hom^ mice would be capable of developing increased amounts of CD8 T cells relative to HLA-B*39:06^Hemi^ mice. We therefore examined the percent of blood CD8 T cells in female NOD.HLA-B*39:06.β2m^KO^ mice (Fig. 2C). HLA-B*39:06^Hemi^ mice had 1.5% CD8 T cells among their peripheral blood leukocytes, while HLA-B*39:06^Hom^ mice had nearly double that amount with 2.5% CD8 T cells, indicating that increased HLA-B*39:06 expression can mediate the development of a higher percentage of CD8 T cells. Mice homozygous for HLA-B*39:06 were used for all subsequent experiments.

**Figure 2.**
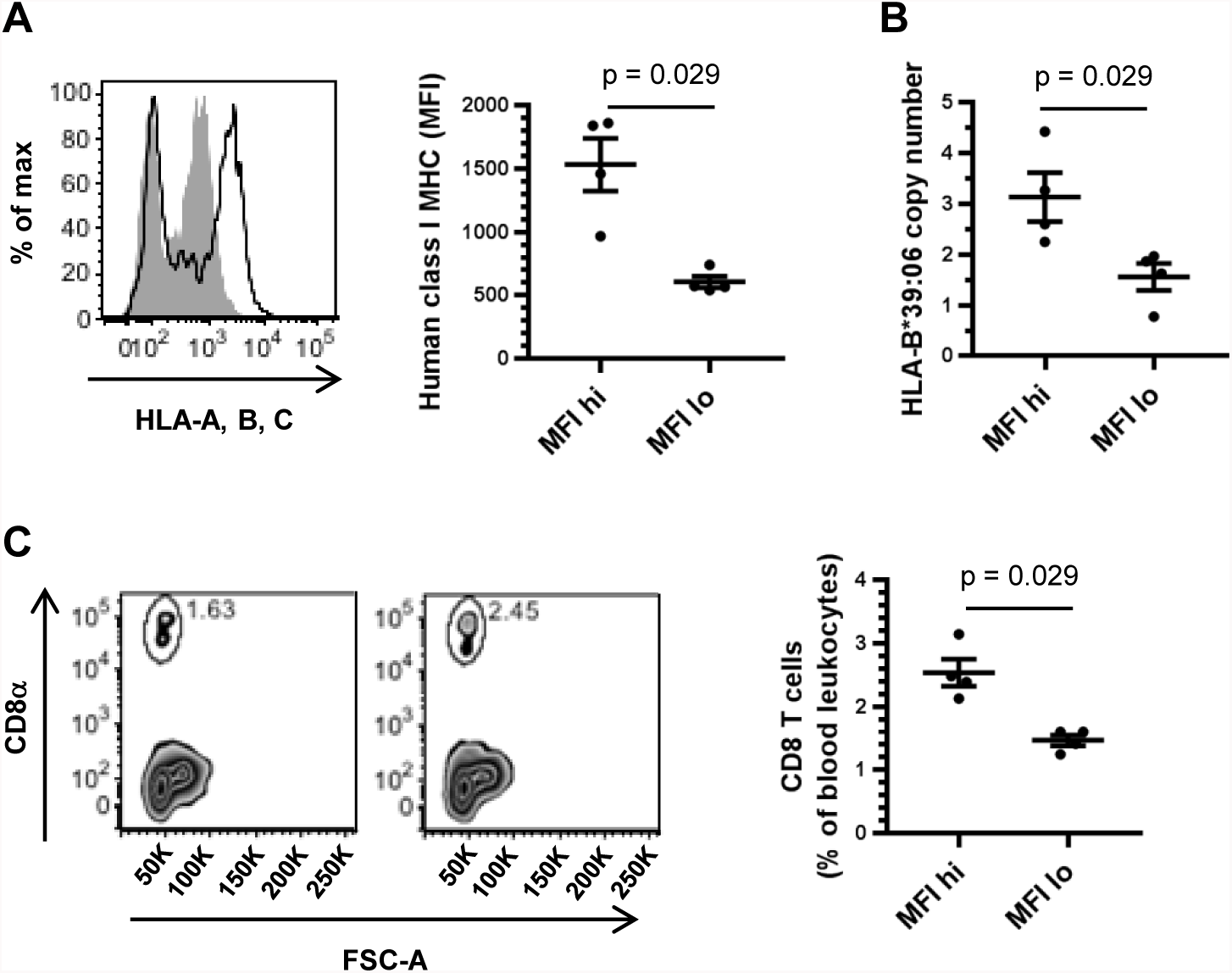
HLA-B*39:06 expression and CD8 T cell development in NOD.HLA-B*39:06.β2m^KO^ hemizygous and homozygous mice. (**A**) Peripheral blood leukocytes from eight NOD.HLA-B*39:06.β2m^KO^ mice were analyzed by flow cytometry for expression of human class I MHC. Left panel: representative histograms for an MFI-high (black line) and an MFI-low mouse (filled gray) are shown. Right panel: geometric MFI of the positive population for each mouse is displayed. Each circle represents an individual mouse. Lines denote mean ± SEM (p = 0.029, Mann-Whitney). (**B**) DNA from four mice per group was assessed for HLA-B*39:06 copy number by quantitative PCR. Each circle represents an individual mouse. Lines denote mean ± SEM (p = 0.029, Mann-Whitney). (**C**) The percent of CD8 T cells among peripheral blood leukocytes was assessed for four mice per group by flow cytometry. Left panels: representative plots for an MFI-low (left) and an MFI-high mouse (right) are shown. Right panel: The percent of CD8 T cells among blood leukocytes is shown. Each circle represents an individual mouse. Lines denote mean ± SEM (p = 0.029, Mann-Whitney).

Having observed CD8 T cells in the peripheral blood of NOD.HLA-B*39:06.β2m^KO^ mice, we next sought to confirm the lack of cell-surface expression of murine class I MHC on splenocytes using the pan murine class I MHC antibody M1/42. Spleens from NOD and NOD.β_2_m^KO^ mice, and the previously characterized NOD.HLA-A*02:01.β2m^KO^ strain (21), were also examined. As expected, only NOD splenocytes showed expression of murine class I MHC (Fig. 3A). The absence of murine class I MHC in NOD.β2m^KO^ mice results in a lack of CD8 T cells (Fig. 3B, 3C), as reported previously (37). However, we observed a partial restoration of CD8 T cell development in the NOD.HLA-B*39:06.β2m^KO^ strain (Fig. 3B, 3C), demonstrating that HLA-B*39:06 is indeed able to mediate CD8 T cell development.

**Figure 3.**
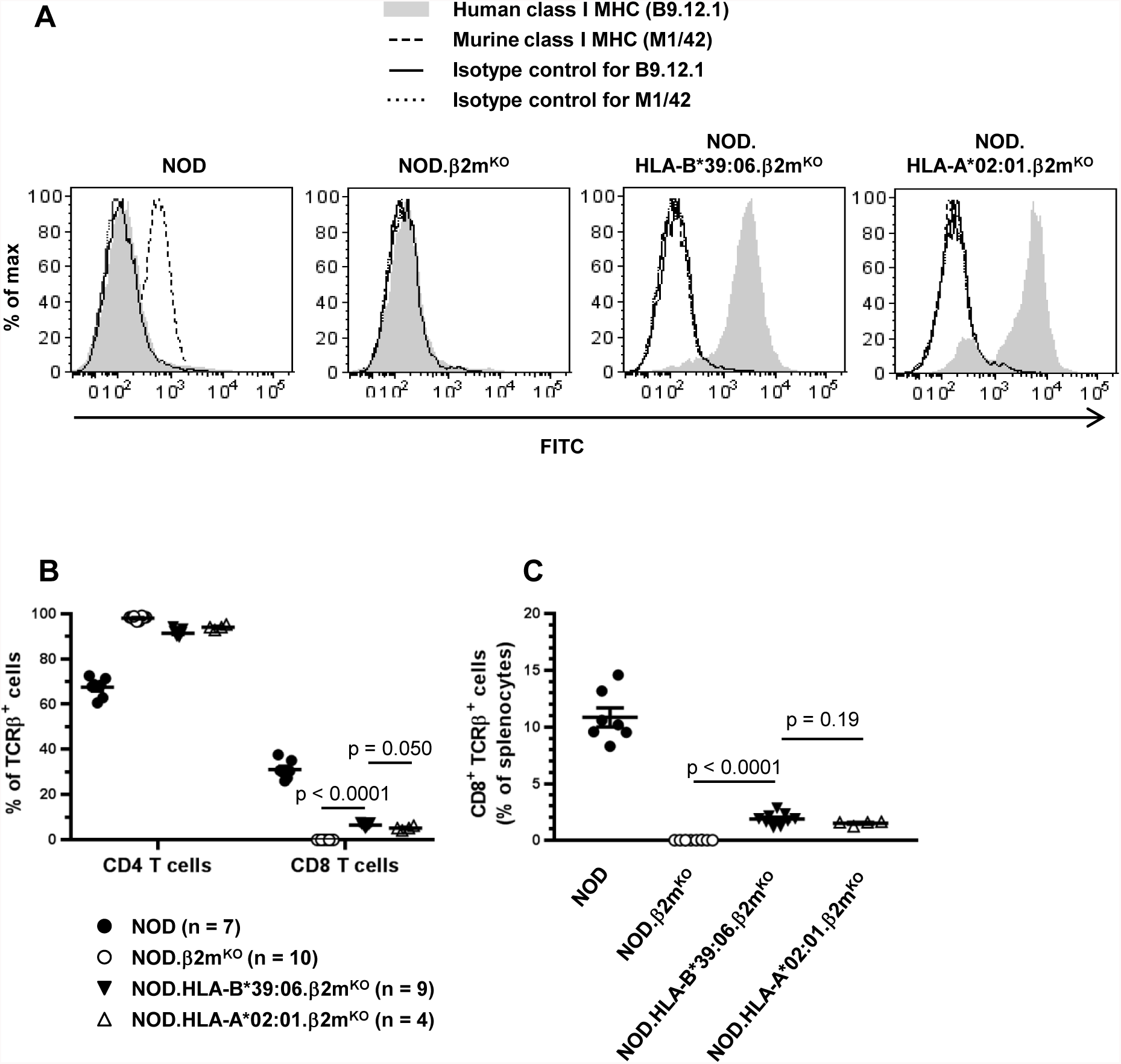
HLA-B*39:06 allows for the development of CD8 T cells in NOD mice. Splenocytes from the indicated mouse strains were analyzed by flow cytometry for human and murine class I MHC expression (**A**) and percentages of the indicated T cell subsets among TCRβ^+^ cells (**B**) or splenocytes (**C**). (B-C) Lines denote mean ± SEM; p values are indicated (Mann-Whitney).

### HLA-B*39:06 mediates T1D in NOD mice lacking murine β2m

We next examined the ability of HLA-B*39:06 to mediate the development of T1D. NOD.β2m^KO^ mice are protected from T1D because they lack CD8 T cells (37, 40-42). However, homozygous expression of HLA-B*39:06 in NOD.β2m^KO^ mice partially restored a disease phenotype (Fig. 4A), with the earliest age of onset at 20 wks and with 17% of NOD.HLA-B*39:06.β2m^KO^ mice diabetic at 40 wks. We therefore show here for the first time that HLA-B*39:06 is able to independently lead to the development of T1D in mice. As previously reported (21), homozygous expression of HLA-A*02:01 (HHD) also allowed for partial restoration of disease susceptibility (Fig. 4A). NOD.HLA-A*02:01.β2m^KO^ mice had their earliest age of onset at 10 wks of age, with 33% diabetic at 40 wks (Fig. 4A). While the age of onset in the HLA-B*39:06 mice was later than that seen in HLA-A*02:01 mice, the two incidence curves were statistically indistinguishable (p = 0.17), and examination of insulitis in n-diabetic mice of each strain revealed similar amounts of islet infiltration (p = 0.15) (Fig. 4B). Representative islets from HLA-A*02:01 and HLA-B*39:06 mice are shown in Fig. 4C and 4D, respectively. Consistent with previous results (27), the majority of NOD.HLA-A*02:01.β2m^KO^ mice displayed insulitis (Fig. 4B). Similarly, histological examination of islets from 40-wk-old NOD.HLA-B*39:06.β2m^KO^ mice revealed that, despite not all progressing to overt T1D, all mice displayed some degree of insulitis, with 81% of mice fully infiltrated (Fig. 4B).

**Figure 4.**
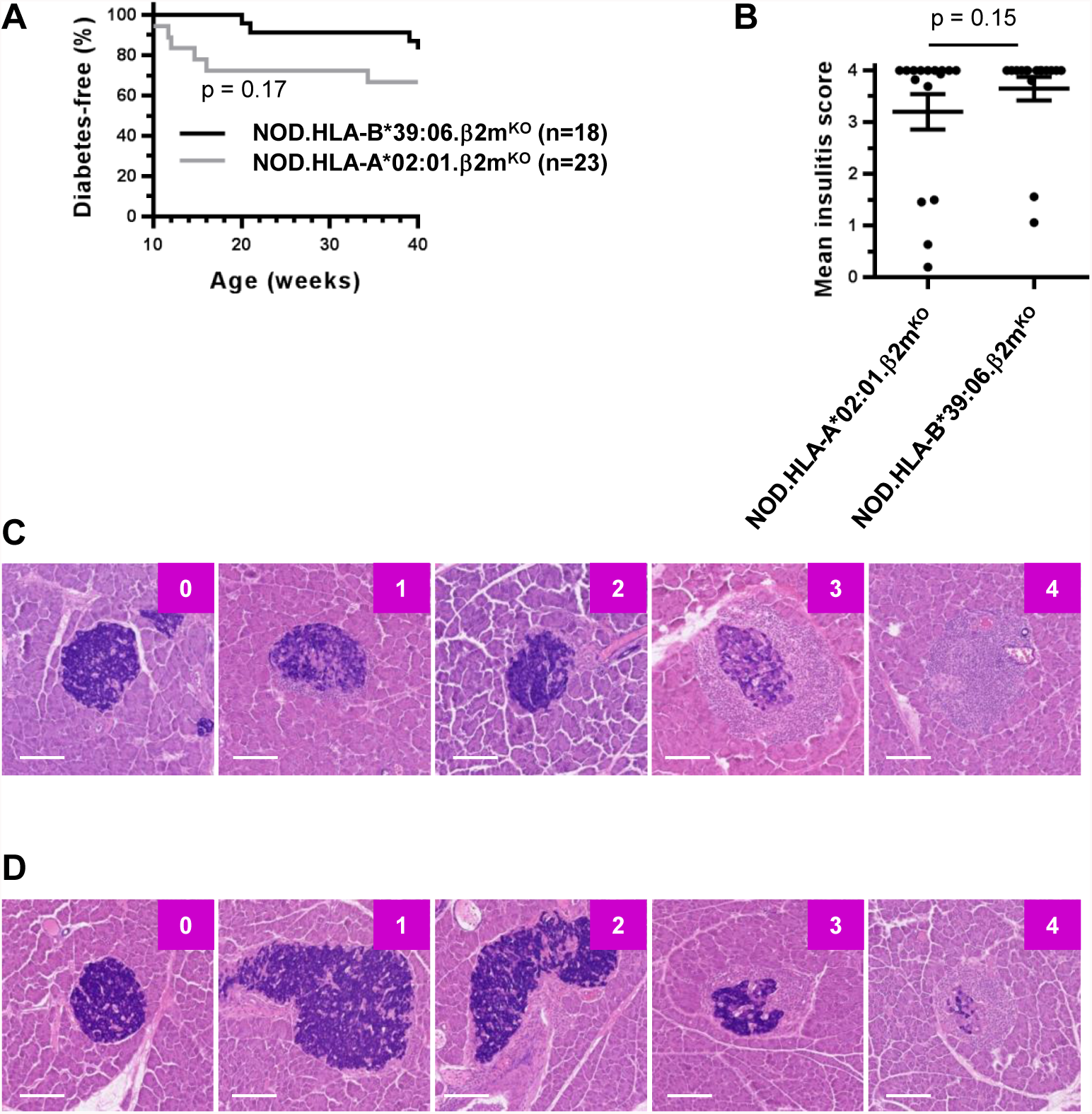
NOD.HLA-B*39:06^Hom^.β2mKO mice are susceptible to T1D. (**A**) The results of diabetes incidence studies performed at The Jackson Laboratory using female NOD.HLA-B*39:06^Hom^.β2m^KO^ and NOD.HLA-A*02:01.β2m^KO^ mice are shown (p = 0.17, Mantel-Cox). (**B**) Female non-diabetic NOD.HLA-A*02:01.β2m^KO^ (n = 16) and NOD.HLA-B*39:06^Hom^.β2m^KO^ mice (n = 16) were euthanized at 40 wks and mean insulitis scores determined as described in *Materials and Methods*. Each circle represents an individual mouse. Lines represent mean ± SEM (p = 0.15, Mann-Whitney). Representative islets with their assigned scores for the female non-diabetic NOD.HLA-A*02:01.β2m^KO^ (**C**) and NOD.HLA-B*39:06^Hom^.β2m^KO^ mice (**D**) are shown. Scale bar represents 100 μm.

### Decreased thymic insulin expression results in earlier T1D onset in HLA-B*39:06-transgenic mice

As little (28, 32) to no (33, 34) *Ins1* expression occurs in the thymus, NOD.Ins2^KO^ mice are characterized by greatly diminished thymic insulin expression (31). They display accelerated T1D onset, increased insulitis, increased T cell reactivity to insulin, and impaired tolerance to insulin compared to WT littermates (29-31). The impact of *Ins2* deficiency on disease is dependent on the genetic context, as NOD.HLA-A*02:01.β2m^KO^.Ins2^KO^ mice have a faster disease onset than NOD.Ins2^KO^ mice (43), indicating that the effects of multiple risk alleles can combine to increase risk. We find that, in conjunction with HLA-B*39:06, *Ins2* deficiency leads to a rapid onset of disease (Fig. 5A, 5B), demonstrating the importance of examining T1D in the context of multiple risk factors. Female NOD.HLA-B*39:06.β2m^KO^ mice had an earliest age of onset of 25 wks, with 58% diabetic at 30 wks. In contrast, NOD.HLA-B*39:06.β2m^KO^.Ins2^KO^ mice had an earliest age of onset of 12 wks, with 100% of this strain being diabetic by 16 wks. We have previously noted that Ins2^Het^ NOD mice exhibit a modest decrease in thymic insulin expression compared to WT NOD mice (27). Comparison of incidence curves (Fig. 5A) and age at onset (Fig. 5B) showed a trend for NOD.HLA-B*39:06.β2m^KO^.Ins2^Het^ mice to exhibit a disease phenotype intermediate between that of their KO and WT counterparts, though the differences between the Het and WT mice did not reach statistical significance with the sample sizes available.

**Figure 5.**
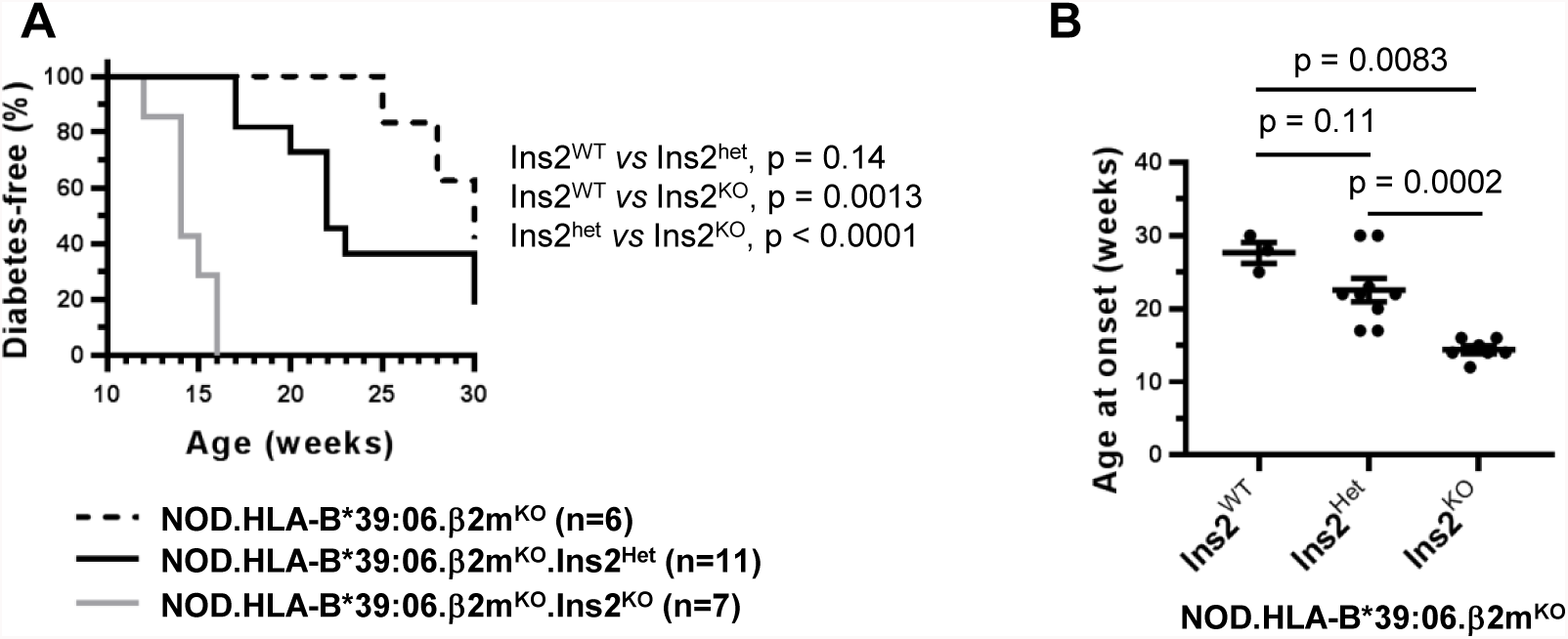
HLA-B*39:06^Hom^ mice display increased diabetes development in the context of reduced thymic insulin expression. (**A**) The results of diabetes incidence studies performed at Albert Einstein College of Medicine using female NOD.HLA-B*39:06^Hom^.β2m^KO^ mice (n = 6) and their Ins^Het^ (n = 11) and Ins2^KO^ (n = 7) counterparts are shown; p values are indicated (Mantel-Cox). (**B**) The ages at onset for all mice in (A) that became diabetic during the incidence study are plotted. Each circle represents an individual mouse. Lines denote mean ± SEM; p values are indicated (Mann-Whitney). Ins2^WT^, n = 3; Ins2^Het^, n = 9; Ins2^KO^, n = 7.

### Differing amounts of thymic insulin expression do not grossly alter lymphocyte populations

To determine whether the increased disease susceptibility seen in the HLA-B*39:06.β2m^KO^.Ins2^KO^ mice was due to gross changes in lymphocyte populations, we examined the impact of differing amounts of thymic insulin expression on splenic B cell, CD8 T cell, and CD4 T cell populations (Fig. 6A). As previously reported, NOD.HLA-A*02:01.β2m^KO^ mice have a reduced percentage of splenic CD8 T cells and an increased percentage of B cells and CD4 T cells relative to NOD mice (21, 23). Given their similar background, we compared our three NOD.HLA-B*39:06.β2m^KO^ strains to age-matched NOD.HLA-A*02:01.β2m^KO^ mice. We found no significant changes in percentage in any of the splenocyte subsets examined (Fig. 6B). The percent of CD8 T cells found in the spleens of HLA-B*39:06 mice was consistent with that found in blood (Fig. 2C). Furthermore, the percentage of CD4^+^CD25^+^ T cells was consistent across all groups, suggesting that the change seen in disease susceptibility was not due to a differing proportion of largely regulatory T cells. Together, these data suggest that the increase in disease incidence seen in the NOD.HLA-B*39:06.β2m^KO^.Ins2^KO^ mice was not due to gross changes in lymphocyte composition compared to the other groups examined.

**Figure 6.**
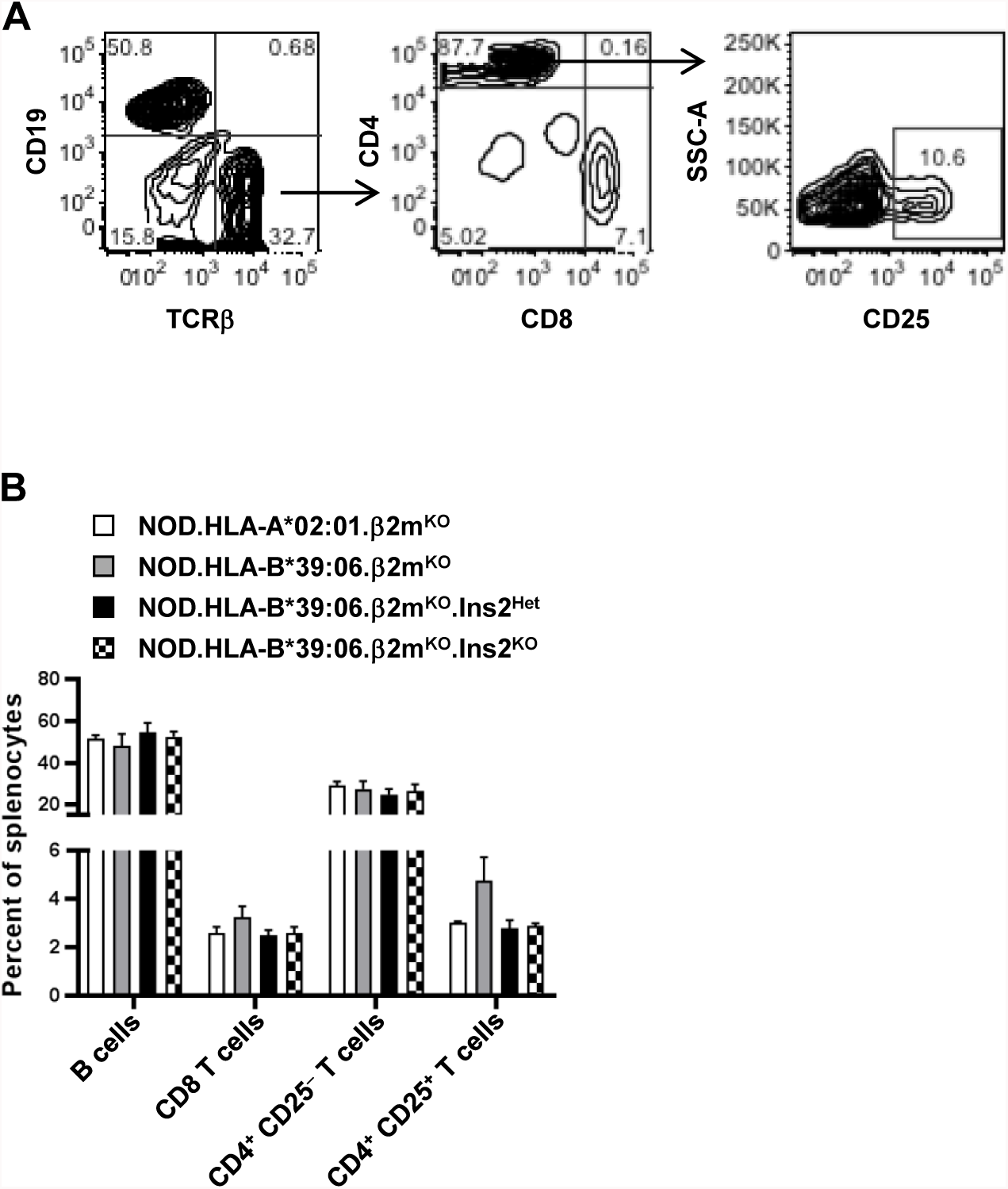
Splenocyte composition does not differ among NOD.HLA-B*39:06^Hom^.β2mKO mice regardless of Ins2 genotype. Splenocytes from three female mice per group (16-25 wks of age) were analyzed by flow cytometry for percentages of lymphocyte populations. (**A**) Gating strategy. (**B**) Percentages of the indicated cell populations are shown. Graph depicts mean + SEM.

### Thymic insulin expression does not alter TCR Vβ usage, but HLA transgene identity does

We next investigated whether changes in TCR Vβ usage accompanied the enhanced disease observed in the NOD.HLA-B*39:06.β2m^KO^.Ins2^KO^ mice. For this purpose, splenocytes from NOD.HLA-B*39:06.β2m^KO^ mice and their Ins2^KO^ counterparts were stained with a panel of anti-mouse TCR Vβ antibodies. Separate examination of CD8 and CD4 T cells revealed no significant differences in TCR Vβ usage between these two strains of mice (Fig. 7A, 7B). When the CD4^+^CD25^+^ (largely regulatory) T cell population was examined individually, the Ins2^KO^ mice showed a small but significant increase in the use of TCR Vβ8.1/2 when compared to Ins2^WT^ mice, but no other changes were noted (Fig. 7C).

**Figure 7.**
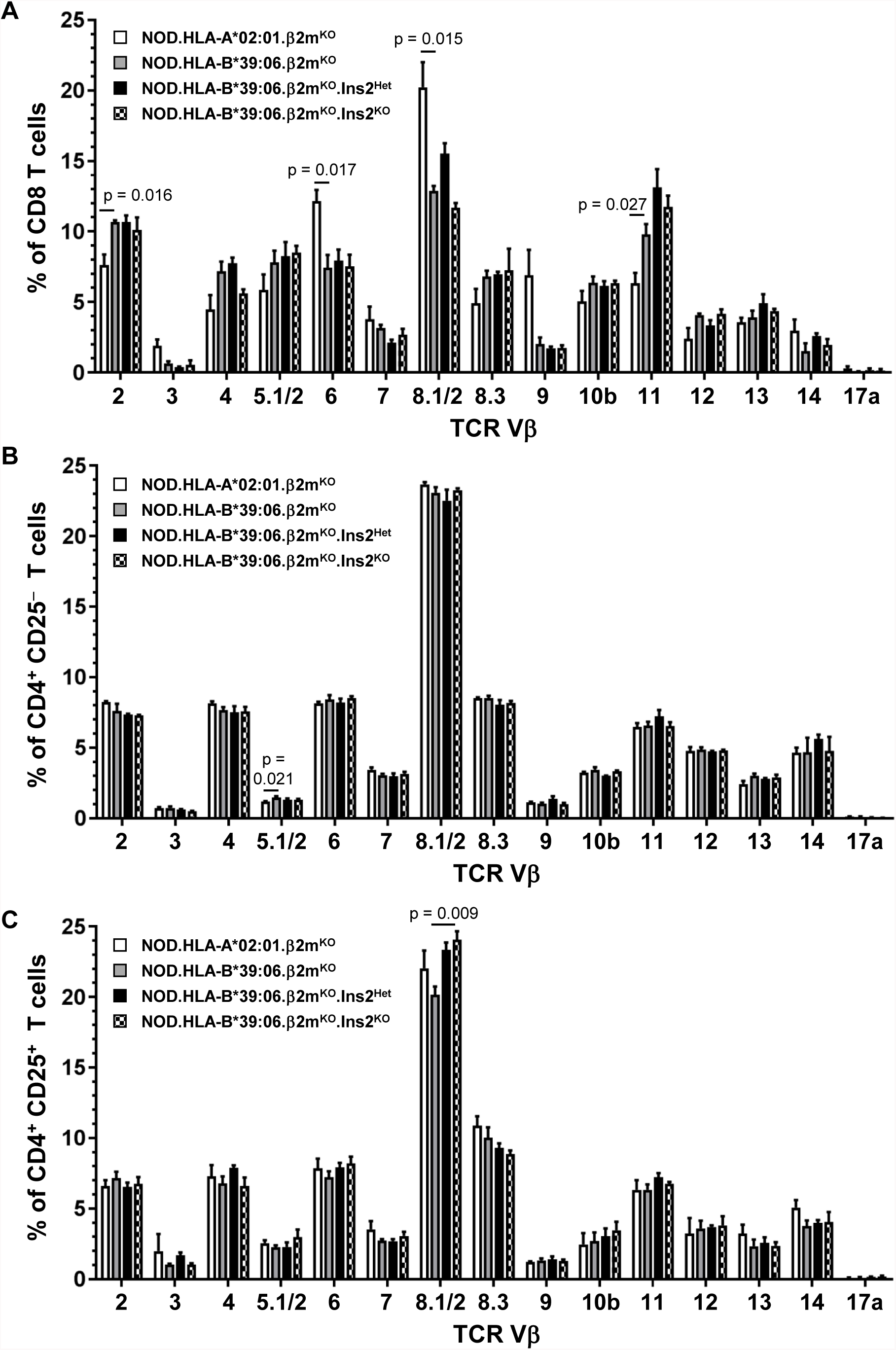
Class I HLA transgene identity dramatically alters TCR Vβ usage among CD8 T cells, but thymic insulin expression does not. Splenocytes from three female mice per group were analyzed by flow cytometry for TCR Vβ usage. (**A**) CD8 T cells, (**B**) CD4 T cells, and (**C**) CD4^+^CD25^+^ T cells were considered separately. Graphs depict mean + SEM; p values are indicated (t test). Significant differences between NOD.HLA-B*39:06.β2m^KO^ and NOD.HLAA*02:01.β2m^KO^ mice are indicated with p values, as are those between NOD.HLA-B*39:06.β2m^KO^ and NOD.HLA-B*39:06.β2m^KO^.Ins2^KO^ mice.

The availability of both NOD.HLA-B*39:06.β2m^KO^ and NOD.HLA-A*02:01.β2m^KO^ mice presented a unique opportunity to examine the influence of HLA transgene identity on TCR Vβ usage. Examination of CD8 T cells revealed significant differences in usage of Vβ2, Vβ6, Vβ8.1/2, and Vβ11 (Fig. 7A). There were no differences in the usage of these TCR Vβ families when CD4 T cell populations were studied (Fig. 7B, 7C), indicating that the presence of HLA-A*02:01 or HLA-B*39:06 specifically alters CD8 T cell selection and/or expansion.

### Despite altered thymic insulin expression, HLA-B*39:06-transgenic NOD mice retain typical blood insulin levels

Because of the compensatory changes observed in pancreatic *Ins* gene expression when the total number of *Ins* genes is reduced from four to two (28), we considered it unlikely that the earlier age of disease onset seen in the NOD.HLA-B*39:06^Hom^.β2m^KO^.Ins2^KO^ mice was due to insufficient pancreatic insulin expression. To confirm this, we measured blood insulin levels in young mice (5-6.5 wks old), well prior to disease onset (Fig. 8). We found that the level of insulin expression was statistically indistinguishable between NOD.HLA-B*39:06^Hom^.β2m^KO^ mice and their Ins2^Het^ and Ins2^KO^ counterparts, with an average concentration of 0.8 ng/ml, consistent with previous reports for other mouse strains (44). This supports the notion that the changes in disease onset are due to diminished immunological tolerance to insulin (30, 31) and not to an inherently decreased ability to produce insulin.

## DISCUSSION

Multiple loci are associated with T1D risk, including a number of class I and class II MHC alleles (4). Among these, HLA-B*39:06 is not only the most predisposing class I HLA allele in T1D patients (7, 9), but also leads to an earlier age of disease onset (11). However, due to the rarity of this allele among the populations studied thus far, investigation of the direct impact of HLA-B*39:06 on T1D pathogenesis has not been possible (6, 7). It is important to note that HLA-B*39:06 is more common among Latin American populations with allele frequencies of 0.03 among Mexican Americans, 0.02-0.06 among Hispanic Americans, and 0.01-0.09 among Mexicans (12). The Venezuela Perja Mountain Bari population has an allele frequency of 0.24. While T1D is relatively rare within Latin American countries, incidence is rising worldwide and new patients from these populations can be expected (14, 15). Similarly, patients carrying this genetic variant can increasingly be found in countries where T1D incidence is highest (45). While genetic background is important, environment is as well; when individuals from areas with low T1D incidence move to areas with high incidence, they assume some of the risk of their new environment (46, 47). Therefore inclusion of HLA-B*39:06-positive patients in treatment studies is essential. As such, the development of a mouse model for the study of HLA-B*39:06 is important as this resource can provide a useful preclinical tool for the testing of HLA-B*39:06-directed treatments in the absence of sufficiently powered patient studies.

We have previously used an NOD.β2m^KO^-based model to study the contribution of HLA-A*02:01 to T1D development (21). In the current study, we found that NOD.HLA-B*39:06^Hom^.β2m^KO^ mice develop similar amounts of CD8 T cells as their HLA-A*02:01-transgenic counterparts (Fig. 3B, 3C), suggesting that HLA-B*39:06 is as efficient at leading to CD8 T cell development as the more well-studied HLA-A*02:01. Indeed, we show here for the first time that HLA-B*39:06 can directly mediate T1D in an NOD.β2m^KO^ model (Fig. 4A, 5A). However, unlike HLA-A*02:01 (21, 48), when expressed in the presence of the NOD class I MHC alleles H2-D^b^ and H2-K^d^, HLA-B*39:06 did not accelerate disease (Fig. 1). While this may be due to strain-specific differences (*e.g.*, transgene integration site), it also may speak to the importance of other aspects of the genetic environment which are known to be important for HLA-B*39:06-related susceptibility in humans. HLA-B*39:06 has been found to exert its effect on T1D risk in patients with specific class II MHC haplotypes, namely HLA-DR8/DQ4 (6, 49). Depending on the population studied, these class II MHCs may be independently predisposing, in which case HLA-B*39:06 accelerates disease progression, or may have a neutral impact on T1D, in which case HLA-B*39:06 lends risk to such patients. It has been well established that H2-A^g7^, the NOD class II MHC, bears great similarity to the human T1D-associated HLA-DQ8 (18). HLA-DR8, part of a haplotype associated with HLA-B*39:06, has similar peptide binding characteristics to both H2-A^g7^ and HLA-DQ8 (50). While HLA-DR8 has been hypothesized to be the T1D-causative allele in the HLA-DR8/DQ4 haplotype, other evidence suggests that HLA-DQ4 is associated with risk of greater disease progression (51-53). Given that genetic context is important for the association between HLA-B*39:06 and T1D risk, the lack of a class II MHC molecule similar to HLA-DQ4 in the NOD mouse model may explain why similar diabetes incidence curves were observed for NOD and NOD.HLA-B*39:06 mice (Fig. 1). However, despite potentially not having an ideal genetic environment, HLA-B*39:06 is still able to mediate disease and islet infiltration, as confirmed by our findings in the NOD.HLA-B*39:06.β2m^KO^ strain (Fig. 4A, 4B, 4D, 5A).

To more accurately model the genetic background of patients with T1D, we incorporated reduced thymic insulin expression into the NOD.HLA-B*39:06.β2m^KO^model. We found that NOD.HLA-B*39:06.β2m^KO^.Ins2^KO^ mice are susceptible to disease at a younger age compared to their Ins2^WT^ counterparts (Fig. 5). Based on our findings that the gross lymphocyte populations (Fig. 6B), TCR Vβ usage (Fig. 7), and blood insulin levels (Fig. 8) do not differ dramatically between these strains, the most likely explanation for the earlier age of onset in the context of reduced insulin expression is a decrease in insulin tolerance (30, 31). CD8 T cells are necessary for the development of T1D (2, 37, 40-42). As the expression of HLA-B*39:06 restores T1D susceptibility to NOD.β2m^KO^ mice (Fig. 4A, 5A) and is enhanced in the Ins2^KO^ mice (Fig. 5A), it is likely that reduced thymic insulin expression results in an increase in CD8 T cell reactivity towards insulin. Increased CD4 T cell reactivity to insulin could also be a contributing factor. We propose that an increased HLA-B*39:06-restricted reactivity to insulin may also contribute to the earlier age of onset seen in HLA-B*39:06-positive patients. These points will be clarified by future investigations.

**Figure 8.**
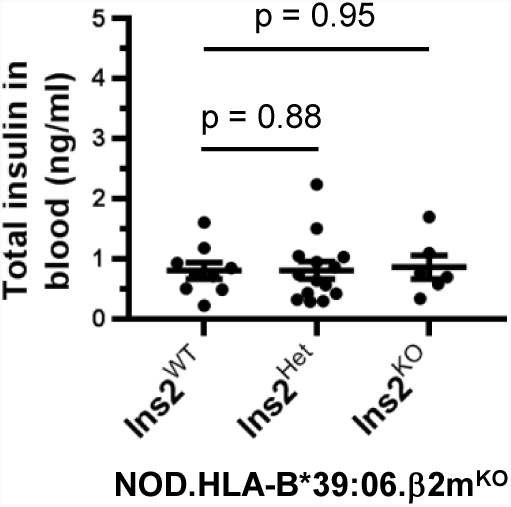
Blood insulin levels in young NOD.HLA-B*39:06.β2m^KO^ mice do not vary regardless of Ins2 genotype. Serum was collected from NOD.HLA-B*39:06^Hom^.β2m^KO^ mice (5-6.5 wks old) having the indicated Ins2 genotypes (Ins2^WT^, n = 9; Ins2^Het^, n = 14; Ins2^KO^, n = 6) in the morning, and insulin levels were measured by ELISA. Each circle represents an individual mouse. Lines denote mean ± SEM; p values are indicated (Mann-Whitney).

The NOD.HLA-B*39:06.β2m^KO^ model can be used in a variety of ways to probe the influence of HLA-B*39:06 on T1D susceptibility. As we have successfully done for HLA-A*02:01 (21, 22, 29), the model will allow us to identify the beta cell peptides recognized by HLA-B*39:06-restricted T cells without the potentially confounding presence of murine class I MHC molecules. That the peptide-binding motif for HLA-B*39:06 has recently been identified may simplify the identification of HLA-B*39:06-restricted epitopes (54, 55). We have previously identified HLA-A*02:01-restricted epitopes in an NOD.β2m^KO^-based model (21, 22, 29); these epitopes were the same or similar to those found in human T1D patients expressing this class I variant (21, 23). Thus the use of the NOD.HLA-B*39:06.β2m^KO^ models could provide a direct translational impact. Identification of HLA-B*39:06-restricted epitopes can allow for their use in epitope-directed therapies; these are an attractive option as they can allow for treatments targeted at specific epitopes without the risk of off-site effects. Furthermore, knowledge of targeted epitopes allows for the tracking of response to therapy, *e.g.*, through the use of peptide-MHC tetramers. Such therapies require preclinical testing, representing another future use of the NOD.HLA-B*39:06.β2m^KO^ models.

Finally, it is important to appreciate that not all HLA class I molecules influence T1D in the same way or to the same degree when expressed in NOD mouse models. For example, when expressed along with H2-D^b^ and H2-K^d^ in NOD mice, HLA-A*02:01 accelerates disease onset (21, 48), HLA-A*11:01 (56) and HLA-B*39:06 (Fig. 1) have no effect, and HLA-B*27 is protective (48). When we compared the TCR Vβ usage among CD8 T cells in mice transgenic for either HLA-A*02:01 or HLA-B*39:06 and lacking murine class I MHC molecules (Fig. 7A), we found that four TCR Vβ families were differentially expressed. This was initially a surprising finding, as until recently it was not generally thought that a given TCR Vβ family had any preference for a particular MHC allele (57). Recently, however, usage of TCR Vβ (and Vα) genes has been found to be associated with MHC genotype in humans, leading to the proposal that different TCR V gene products may indeed have a bias toward particular MHC alleles (58). Our results using the NOD.HLA-B*39:06.β2m^KO^ and NOD.HLA-A*02:01.β2m^KO^ strains (Fig. 7A) support this view and represent a valuable system to explore this phenomenon further. Differences in TCR repertoire could help to explain the differential T1D susceptibility observed not only in NOD mice transgenic for different HLA class I alleles, but, more importantly, in humans as well.

In sum, we have established that HLA-B*39:06 can directly mediate T1D in the NOD mouse model, confirming the results seen in multiple genome-wide association studies (5, 7). We have furthermore developed models for HLA-B*39:06 in a genetic context more relevant to human disease by incorporating reduced thymic insulin expression. These models will allow a detailed investigation of the influence of HLA-B*39:06 on T1D development.

## ACKNOWLEDGMENTS

We thank Denisa Ferastraoaru for assistance with the preparation of the monochain chimeric HLA-B*39:06 construct. We acknowledge The Jackson Laboratory Transgenic Genotyping Core for developing the real-time PCR assay for monitoring HLA-B*39:06 homozygosity.

## FOOTNOTES

2 Address correspondence and reprint requests to Teresa P. DiLorenzo (Department of Microbiology and Immunology, Albert Einstein College of Medicine, 1300 Morris Park Ave., Bronx, NY 10461; E-mail address)

3 Abbreviations used in this paper: β2m, β_2_-microglobulin; Het, heterozygous; HHD, human β2m/HLA/H2-D^b^; Hom, homozygous; Ins2, insulin 2; KO, knockout; MFI, mean fluorescence intensity; T1D, type 1 diabetes; VNTR, variable number of tandem repeats; WT, wild-type.

## REFERENCES

1. Roep, B. O., and M. Peakman. 2011. Diabetogenic T lymphocytes in human Type 1 diabetes. Curr. Opin. Immunol. 23: 746–753.

2. Varela-Calvino, R., C. Calvino-Sampedro, I. Gomez-Tourino, and O. J. Cordero. 2017. Apportioning blame: autoreactive CD4^+^ and CD8^+^ T cells in type 1 diabetes. Arch. Immunol. Ther. Exp. (Warsz.) 65: 275–284.

3. Ferris, S. T., J. A. Carrero, and E. R. Unanue. 2016. Antigen presentation events during the initiation of autoimmune diabetes in the NOD mouse. J. Autoimmun. 71: 19–25.

4. Noble, J. A., and A. M. Valdes. 2011. Genetics of the HLA region in the prediction of type 1 diabetes. Curr. Diab. Rep. 11: 533–542.

5. Howson, J. M., N. M. Walker, D. Clayton, and J. A. Todd. 2009. Confirmation of HLA class II independent type 1 diabetes associations in the major histocompatibility complex including HLA-B and HLA-A. Diabetes Obes. Metab. 11 Suppl 1: 31–45.

6. Mikk, M. L., T. Heikkinen, M. I. El-Amir, M. Kiviniemi, A. P. Laine, T. Harkonen, R. Veijola, J. Toppari, M. Knip, and J. Ilonen. 2017. The association of the HLA-A*24:02, B*39:01 and B*39:06 alleles with type 1 diabetes is restricted to specific HLA-DR/DQ haplotypes in Finns. HLA 89: 215–224.

7. Nejentsev, S., J. M. Howson, N. M. Walker, J. Szeszko, S. F. Field, H. E. Stevens, P. Reynolds, M. Hardy, E. King, J. Masters, J. Hulme, L. M. Maier, D. Smyth, R. Bailey, J. D. Cooper, G. Ribas, R. D. Campbell, D. G. Clayton, and J. A. Todd. 2007. Localization of type 1 diabetes susceptibility to the MHC class I genes HLA-B and HLA-A. Nature 450: 887–892.

8. Noble, J. A., A. M. Valdes, T. L. Bugawan, R. J. Apple, G. Thomson, and H. A. Erlich. 2002. The HLA class I A locus affects susceptibility to type 1 diabetes. Hum. Immunol. 63: 657–664.

9. Noble, J. A., A. M. Valdes, M. D. Varney, J. A. Carlson, P. Moonsamy, A. L. Fear, J. A. Lane, E. Lavant, R. Rappner, A. Louey, P. Concannon, J. C. Mychaleckyj, and H. A. Erlich. 2010. HLA class I and genetic susceptibility to type 1 diabetes: results from the Type 1 Diabetes Genetics Consortium. Diabetes 59: 2972–2979.

10. DiLorenzo, T. P., M. Peakman, and B. O. Roep. 2007. Translational mini-review series on type 1 diabetes: Systematic analysis of T cell epitopes in autoimmune diabetes. Clin. Exp. Immunol. 148: 1–16.

11. Valdes, A. M., H. A. Erlich, and J. A. Noble. 2005. Human leukocyte antigen class I B and C loci contribute to Type 1 Diabetes (T1D) susceptibility and age at T1D onset. Hum. Immunol. 66: 301–313.

12. Gonzalez-Galarza, F. F., L. Y. Takeshita, E. J. Santos, F. Kempson, M. H. Maia, A. L. da Silva, A. L. Teles e Silva, G. S. Ghattaoraya, A. Alfirevic, A. R. Jones, and D. Middleton. 2015. Allele Frequency Net 2015 update: new features for HLA epitopes, KIR and disease and HLA adverse drug reaction associations. Nucleic Acids Res. 43: D784–788.

13. Lerner, A., P. Jeremias, and T. Matthias. 2015. The world incidence and prevalence of autoimmune diseases is increasing. Int. J. Celiac Dis. 3: 151–155.

14. Carrasco, E., F. Perez-Bravo, J. Dorman, A. Mondragon, and J. L. Santos. 2006. Increasing incidence of type 1 diabetes in population from Santiago of Chile: trends in a period of 18 years (1986-2003). Diabetes Metab. Res. Rev. 22: 34–37.

15. Gomez-Diaz, R. A., N. Garibay-Nieto, N. Wacher-Rodarte, and C. A. Aguilar-Salinas. 2014. Epidemiology of type 1 diabetes in Latin America. Curr. Diabetes Rev. 10: 75–85.

16. Anderson, M. S., and J. A. Bluestone. 2005. The NOD mouse: a model of immune dysregulation. Annu. Rev. Immunol. 23: 447–485.

17. Driver, J. P., Y. G. Chen, and C. E. Mathews. 2012. Comparative genetics: synergizing human and NOD mouse studies for identifying genetic causation of type 1 diabetes. Rev. Diabet. Stud. 9: 169–187.

18. Suri, A., and E. R. Unanue. 2005. The murine diabetogenic class II histocompatibility molecule I-A^g7^: structural and functional properties and specificity of peptide selection. Adv. Immunol. 88: 235–265.

19. Askenasy, N. 2016. Mechanisms of autoimmunity in the non-obese diabetic mouse: effector/regulatory cell equilibrium during peak inflammation. Immunology 147: 377–388.

20. Bettini, M., and D. A. Vignali. 2009. Regulatory T cells and inhibitory cytokines in autoimmunity. Curr. Opin. Immunol. 21: 612–618.

21. Takaki, T., M. P. Marron, C. E. Mathews, S. T. Guttmann, R. Bottino, M. Trucco, T. P. DiLorenzo, and D. V. Serreze. 2006. HLA-A*0201-restricted T cells from humanized NOD mice recognize autoantigens of potential clinical relevance to type 1 diabetes. J. Immunol. 176: 3257–3265.

22. Jarchum, I., J. C. Baker, T. Yamada, T. Takaki, M. P. Marron, D. V. Serreze, and T. P. DiLorenzo. 2007. In vivo cytotoxicity of insulin-specific CD8^+^ T-cells in HLA-A*0201 transgenic NOD mice. Diabetes 56: 2551–2560.

23. Jarchum, I., L. Nichol, M. Trucco, P. Santamaria, and T. P. DiLorenzo. 2008. Identification of novel IGRP epitopes targeted in type 1 diabetes patients. Clin. Immunol. 127: 359–365.

24. Vafiadis, P., S. T. Bennett, J. A. Todd, J. Nadeau, R. Grabs, C. G. Goodyer, S. Wickramasinghe, E. Colle, and C. Polychronakos. 1997. Insulin expression in human thymus is modulated by INS VNTR alleles at the IDDM2 locus. Nat. Genet. 15: 289–292.

25. Kelly, M. A., M. L. Rayner, C. H. Mijovic, and A. H. Barnett. 2003. Molecular aspects of type 1 diabetes. Mol. Pathol. 56: 1–10.

26. Howson, J. M., N. M. Walker, D. J. Smyth, and J. A. Todd. 2009. Analysis of 19 genes for association with type I diabetes in the Type I Diabetes Genetics Consortium families. Genes Immun. 10 Suppl 1: S74–84.

27. Babad, J., R. Ali, J. Schloss, and T. P. DiLorenzo. 2016. An HLA-transgenic mouse model of type 1 diabetes that incorporates the reduced but not abolished thymic insulin expression seen in patients. J. Diabetes Res. 2016: 7959060.

28. Chentoufi, A. A., and C. Polychronakos. 2002. Insulin expression levels in the thymus modulate insulin-specific autoreactive T-cell tolerance: the mechanism by which the *IDDM2* locus may predispose to diabetes. Diabetes 51: 1383–1390.

29. Jarchum, I., and T. P. DiLorenzo. 2010. Ins2 deficiency augments spontaneous HLA-A*0201-restricted T cell responses to insulin. J. Immunol. 184: 658–665.

30. Moriyama, H., N. Abiru, J. Paronen, K. Sikora, E. Liu, D. Miao, D. Devendra, J. Beilke, R. Gianani, R. G. Gill, and G. S. Eisenbarth. 2003. Evidence for a primary islet autoantigen (preproinsulin 1) for insulitis and diabetes in the nonobese diabetic mouse. Proc. Natl. Acad. Sci. U. S. A. 100: 10376–10381.

31. Thebault-Baumont, K., D. Dubois-Laforgue, P. Krief, J. P. Briand, P. Halbout, K. Vallon-Geoffroy, J. Morin, V. Laloux, A. Lehuen, J. C. Carel, J. Jami, S. Muller, and C. Boitard. 2003. Acceleration of type 1 diabetes mellitus in proinsulin 2-deficient NOD mice. J. Clin. Invest. 111: 851–857.

32. Fan, Y., W. A. Rudert, M. Grupillo, J. He, G. Sisino, and M. Trucco. 2009. Thymus-specific deletion of insulin induces autoimmune diabetes. EMBO J. 28: 2812–2824.

33. Deltour, L., P. Leduque, N. Blume, O. Madsen, P. Dubois, J. Jami, and D. Bucchini. 1993. Differential expression of the two nonallelic proinsulin genes in the developing mouse embryo. Proc. Natl. Acad. Sci. U. S. A. 90: 527–531.

34. Heath, V. L., N. C. Moore, S. M. Parnell, and D. W. Mason. 1998. Intrathymic expression of genes involved in organ specific autoimmune disease. J. Autoimmun. 11: 309–318.

35. Faideau, B., J. P. Briand, C. Lotton, I. Tardivel, P. Halbout, J. Jami, J. F. Elliott, P. Krief, S. Muller, C. Boitard, and J. C. Carel. 2004. Expression of preproinsulin-2 gene shapes the immune response to preproinsulin in normal mice. J. Immunol. 172: 25–33.

36. Pascolo, S., N. Bervas, J. M. Ure, A. G. Smith, F. A. Lemonnier, and B. Perarnau. 1997. HLA-A2.1-restricted education and cytolytic activity of CD8^+^ T lymphocytes from β2 microglobulin (β2m) HLA-A2.1 monochain transgenic H-2D^b^ β2m double knockout mice. J. Exp. Med. 185: 2043–2051.

37. Serreze, D. V., E. H. Leiter, G. J. Christianson, D. Greiner, and D. C. Roopenian. 1994. Major histocompatibility complex class I-deficient NOD-*B2m^null^* mice are diabetes and insulitis resistant. Diabetes 43: 505–509.

38. Cooper, J. C., G. B. Dealtry, M. A. Ahmed, P. C. Arck, B. F. Klapp, S. M. Blois, and N. Fernandez. 2007. An impaired breeding phenotype in mice with a genetic deletion of beta-2 microglobulin and diminished MHC class I expression: role in reproductive fitness. Biol. Reprod. 77: 274–279.

39. Serreze, D. V., H. D. Chapman, D. S. Varnum, M. S. Hanson, P. C. Reifsnyder, S. D. Richard, S. A. Fleming, E. H. Leiter, and L. D. Shultz. 1996. B lymphocytes are essential for the initiation of T cell-mediated autoimmune diabetes: analysis of a new “speed congenic” stock of NOD. *Igμ^null^* mice. J. Exp. Med. 184: 2049–2053.

40. Sumida, T., M. Furukawa, A. Sakamoto, T. Namekawa, T. Maeda, M. Zijlstra, I. Iwamoto, T. Koike, S. Yoshida, H. Tomioka, and et al. 1994. Prevention of insulitis and diabetes in β_2_-microglobulin-deficient non-obese diabetic mice. Int. Immunol. 6: 1445–1449.

41. Wicker, L. S., E. H. Leiter, J. A. Todd, R. J. Renjilian, E. Peterson, P. A. Fischer, P. L. Podolin, M. Zijlstra, R. Jaenisch, and L. B. Peterson. 1994. β2-microglobulin-deficient NOD mice do not develop insulitis or diabetes. Diabetes 43: 500–504.

42. Katz, J., C. Benoist, and D. Mathis. 1993. Major histocompatibility complex class I molecules are required for the development of insulitis in non-obese diabetic mice. Eur. J. Immunol. 23: 3358–3360.

43. Jarchum, I., and T. P. DiLorenzo. 2010. Ins2 deficiency augments spontaneous HLA-A*0201-restricted T cell responses to insulin. Journal of immunology (Baltimore, Md.: 1950) 184: 658–665.

44. Coulaud, J., S. Durant, and F. Homo-Delarche. 2010. Glucose homeostasis in pre-diabetic NOD and lymphocyte-deficient NOD/SCID mice during gestation. Rev. Diabet. Stud. 7: 36–46.

45. Hsin, O., A. M. La Greca, J. Valenzuela, C. T. Moine, and A. Delamater. 2010. Adherence and glycemic control among Hispanic youth with type 1 diabetes: role of family involvement and acculturation. J. Pediatr. Psychol. 35: 156–166.

46. Soderstrom, U., J. Aman, and A. Hjern. 2012. Being born in Sweden increases the risk for type 1 diabetes - a study of migration of children to Sweden as a natural experiment. Acta Paediatr. 101: 73–77.

47. Hussen, H. I., D. Yang, S. Cnattingius, and T. Moradi. 2013. Type I diabetes among children and young adults: the role of country of birth, socioeconomic position and sex. Pediatr. Diabetes 14: 138–148.

48. Marron, M. P., R. T. Graser, H. D. Chapman, and D. V. Serreze. 2002. Functional evidence for the mediation of diabetogenic T cell responses by HLA-A2.1 MHC class I molecules through transgenic expression in NOD mice. Proc. Natl. Acad. Sci. U. S. A. 99: 13753–13758.

49. Baschal, E. E., P. R. Baker, K. R. Eyring, J. C. Siebert, J. M. Jasinski, and G. S. Eisenbarth. 2011. The HLA-B 3906 allele imparts a high risk of diabetes only on specific HLA-DR/DQ haplotypes. Diabetologia 54: 1702–1709.

50. Muixi, L., M. Gay, P. M. Munoz-Torres, C. Guitart, J. Cedano, J. Abian, I. Alvarez, and D. Jaraquemada. 2011. The peptide-binding motif of HLA-DR8 shares important structural features with other type 1 diabetes-associated alleles. Genes Immun. 12: 504–512.

51. Mimura, T., H. Funatsu, Y. Uchigata, S. Kitano, H. Noma, E. Shimizu, Y. Konno, S. Amano, M. Araie, O. Yoshino, Y. Iwamoto, and S. Hori. 2003. Relationship between human leukocyte antigen status and proliferative diabetic retinopathy in patients with younger-onset type 1 diabetes mellitus. Am. J. Ophthalmol. 135: 844–848.

52. Hanafusa, T., and A. Imagawa. 2007. Fulminant type 1 diabetes: a novel clinical entity requiring special attention by all medical practitioners. Nat. Clin. Pract. Endocrinol Metab. 3: 36–45.

53. Parry, C. S., and B. R. Brooks. 2008. A new model defines the minimal set of polymorphism in HLA-DQ and-DR that determines susceptibility and resistance to autoimmune diabetes. Biol. Direct 3: 42.

54. Eichmann, M., A. de Ru, P. A. van Veelen, M. Peakman, and D. Kronenberg-Versteeg. 2014. Identification and characterisation of peptide binding motifs of six autoimmune disease-associated human leukocyte antigen-class I molecules including HLA-B*39:06. Tissue Antigens 84: 378–388.

55. Sidney, J., J. Schloss, C. Moore, M. Lindvall, A. Wriston, D. F. Hunt, J. Shabanowitz, T. P. DiLorenzo, and A. Sette. 2016. Characterization of the peptide binding specificity of the HLA class I alleles B*38:01 and B*39:06. Immunogenetics 68: 231–236.

56. Antal, Z., J. C. Baker, C. Smith, I. Jarchum, J. Babad, G. Mukherjee, Y. Yang, J. Sidney, A. Sette, P. Santamaria, and T. P. DiLorenzo. 2012. Beyond HLA-A*0201: New HLA-transgenic NOD mouse models of type 1 diabetes identify the insulin C-peptide as a rich source of CD8^+^ T cell epitopes. J. Immunol. 188: 5766–5775.

57. Garcia, K. C., J. J. Adams, D. Feng, and L. K. Ely. 2009. The molecular basis of TCR germline bias for MHC is surprisingly simple. Nat. Immunol. 10: 143–147.

58. Sharon, E., L. V. Sibener, A. Battle, H. B. Fraser, K. C. Garcia, and J. K. Pritchard. 2016. Genetic variation in MHC proteins is associated with T cell receptor expression biases. Nat. Genet. 48: 995–1002.

